# Temporal dynamics improves machine learning-based prediction of cell state from quantitative phase imaging

**DOI:** 10.64898/2026.08.24.746855

**Authors:** Shukran Alizada, Kayla A. Marks, Rebecca G. Zitnay, Anne Done, Robert L. Judson-Torres, Thomas A. Zangle

## Abstract

Cell morphology reflects cell health and can distinguish cell-cycle stage, growth arrest, and distinct pathways of cell death. Live, label-free quantitative phase imaging (QPI) captures these features non-invasively and with high temporal resolution, yet many image-based classifiers rely on single frames and cannot separate states whose differences emerge only over time. How much temporal information is needed, and which architecture best exploits it, remain open questions. We assembled 1,874 QPI timelapse sequences spanning six cell states (interphase, mitosis, cell cycle arrest, apoptosis, ferroptosis, and necroptosis) and compared two-dimensional convolutional neural networks (CNNs) with a three-dimensional (3D) spatiotemporal CNN across increasing frame counts. Accuracy improved as frames were added, with the largest gain between one and three frames. The 2D models saturated beyond three frames, whereas the 3D architecture kept improving, reaching 96.5% accuracy and a 3.5% error rate at eleven frames. The temporal information needed tracked the timescale of each process: mitosis was resolved from a single frame, while ferroptosis benefited most from extended sequences. Overall, these results show that dynamic information, rather than static morphology alone, drives accurate cell-state classification, and that 3D architectures are needed to fully exploit it for label-free dynamic phenotyping.

## Introduction

Measurements of cell proliferation, growth dynamics, viability, and phenotypic changes are essential for understanding the mechanisms underlying cell and cancer biology. This need is reflected in the wide array of specialized assays developed for these purposes, such as microplate assays, flow cytometry assays, and high-throughput imaging assays (1,2). When interpreted independently, many standard assays for evaluating cell behavior are ambiguous. For example, distinguishing cytostatic from cytotoxic effects of pharmacological compounds often requires multiple complementary assays (3,4). Multi-parametric measurements of cell morphology derived from image-based single-cell phenotypic profiling can distinguish cellular behaviors, such as growth arrest from distinct cell death pathways, while providing accurate and scalable analysis, leading to mechanistic insight, enabling detection of rare events such as emergent resistance phenotypes, and supporting patient-specific therapeutic decision-making at a fraction of the cost of multi-assay approaches (5–9).

High-content imaging of fixed, labeled cells has highlighted the utility of image-profiling based approaches for cell classification (10), including evaluation of cell health (8), drug discovery (11), and discerning mechanisms of cell death (12). However, approaches based on fixed cell imaging capture only a single predetermined snapshot of inherently dynamic processes. Live imaging, on the other hand, captures cells that are actively cycling, changing state, or dying, providing a lens into cellular processes that are unavailable in a single image. However, many approaches to classify cell processes have used single frames for classification, which can make it difficult to distinguish subtly different states (13–17).

One solution to address this problem is to encode the dynamic data as a feature trajectory which has been used to classify cellular changes, such as the mechanism of cell death (18), drug mechanism of action (19), cell differentiation state (20) or type of cell motility (21). Alternatively, time-series data can be input in a manner that captures the dynamic changes in an image format, such as an image that represents the pixel intensity change from one frame to the next (22), or as a multi-frame clip (23). Evaluating time-series data with machine learning tools can illuminate how cell phenotype changes over time, revealing previously unknown intermediate states, and highlighting morphological features associated with a distinct cell behavior (18,20).

Quantitative phase imaging (QPI), which quantitatively maps a sample’s mass by measuring the phase shift of light through the sample, results in a multi-dimensional information-rich feature set that can be used to image dynamic cell phenotypes (24,25). This technique is especially powerful for live imaging because it does not require adding dyes or invasive materials into cells and uses low-powered light sources that have minimal cytotoxic effects, enabling long-duration imaging with high temporal resolution. Its utility in image profiling has already been well validated across a variety of different QPI platforms including holography (5,7,23).

One advantage of QPI is that it produces strong contrast between biological cells and the surrounding medium by measuring variations in optical phase caused by differences in refractive index and specimen thickness. As a result, cell boundaries are often much easier to distinguish from the background than in conventional bright-field images (24,26). This quantitative information allows image-processing software to apply segmentation methods such as thresholding or, more recently, deep learning methods such as convolutional neural networks (CNNs) and U-Net architectures. Artificial intelligence has become important because it can accurately segment touching or overlapping cells that are difficult to separate using traditional image-processing techniques (27). After segmentation, each detected cell is assigned a unique object identification. The software calculates the boundary or bounding box surrounding that individual cell and crops it from the original image. This creates a separate image containing only that object. Cropping individual objects has several advantages. It allows each cell to be analyzed independently, reduces computational cost by removing unnecessary background information, and provides standardized inputs for machine learning algorithms. The cropped cell images can then be used for classification, tracking over time, morphology measurements, or disease diagnosis. Modern AI-enabled QPI workflows commonly perform segmentation first, followed by object identification and cropping before classification or quantitative analysis (27,28).

Live cell imaging via QPI can be used for identification of cell states. However, how much information is required to identify subtly different cell states remains a key question. Inclusion of additional frames can likely improve identification of dynamic processes, but at the expense of computational and storage costs. Here, we investigate the impact of including temporal information from QPI on cell state classification for 1,874 QPI timelapse sequences representing six distinct cell states: apoptosis, cell cycle arrest, ferroptosis, interphase, mitosis, and necroptosis. First, we performed a broad 2D convolutional neural network (CNN) architecture search to identify the best-performing 2D model, followed by a temporal scaling study looking at state-specific performance across increasing frame counts. We also compare the performance of including time in a 2D CNN to a purpose-built 3D spatiotemporal CNN. Overall, our results show that performance for classification of dynamic cell states improves with the inclusion of additional frames in a manner dependent on phenotype. These results can be used to guide future development of cell state classification methods.

## Materials and Methods

### Cell Culture

Human melanoma cell line, 624-Mel, was maintained in RPMI media containing 10% FBS (Fetal Bovine Serum), 1% penicillin-streptomycin, and 1% NEAA (Non-Essential Amino Acid). 624-Mel cells were grown in the incubator at 37°C and 5% CO2. Cells were passaged using trypsin for dissociation every 3 days at a 1:10 subculture ratio.

### Drug Treatments

The day before imaging, 60,000 cells per well are plated in a 6 well Sarstedt TC Plate in 3ml RPMI. Immediately prior to imaging, media was replaced with 3 mL fresh, warmed RPMI containing treatment compounds at the following concentrations: Staurosporine: 0.001uM, 0.01uM, 0.1uM, 10uM; Shikonin: 0.1uM, 0.5uM, 1uM, 5uM, 10uM; Erastin: 0.1uM, 0.5uM, 1uM, 10uM, 30uM; Dabrafenib: 0.1nM, 1nM, 10nM, 100nM, 1uM; Palbociclib: 0.5nM, 5nM, 50nM, 500nM, 5uM; Barasertib: 0.1nM, 1nM, 10nM, 100nM, 300nM.

### QPI imaging

QPI was performed on the M4 HoloMonitor (Phase Holographic Imaging, Lund, Sweden). Prior to imaging a HoloLid for 6-well plates (Phase Holographic Imaging, Lund, Sweden) was cleaned with a Q-Tip in 70% ethanol. Cells were imaged for 48 hours, with images taken at 15-minute intervals at 12 random imaging locations and 20x magnification in a humidified incubator at 37°C and 5% CO2.

### Image selection

M4 images were exported as 16 bit TIFF images and imported into Fiji. Cell coordinates were determined by identifying the center of the cell in the frame of interest, and manually recording the x, y, and z (time), coordinates. Each cell was identified in a event-specific manner. For cell death phenotypes, apoptosis, necroptosis, and ferroptosis, T0 is determined as the frame prior to obvious cell death, with T+1 being the cell demonstrating features consistent with death. For apoptotic cells, T0 is defined as condensation of the nucleus and cytoplasm, observed as rounding up of the cell. T+1 showing all cell dendrites released from the plates and detachment indicative of death. The identification of necroptotic cells is defined as the rounding up of the cell before it dies. Ferroptotic cells T0 was identified as the frame before a major drop in cell mass. T+1 was defined as when the cell as lost all mass and visually looks transparent and porous. Necroptotic cells T0 were defined as the frame before the rounding up of the cell and before the condensation of the cell shape. T+1 was defines as the cell becoming balled up before death and no dendrites. G0/G1 arrest and G2/M arrest T0 was defined by no cell division for 25-30 hours form the last mitotic event. Late interphase T0 were identified as 6 hours after or 12 hours before a mitotic event, whereas early interphase was 6 hours before or 12 hours after a mitotic event. Mitosis T0 was defined as cell rounding up and T+1 shows two daughter cells.

### Flow procedures

60,000 cells per well are plated in a 6 well plate in 3ml RPMI. The next day cells were treated with one of 5 different small molecules at the concentrations previously reported and one DMSO control, with same treatment procedure as M4 imaging.

For Annexin V, Cells were centrifuged for 5 minutes at 500g, supernatant was removed, and cells are resuspended in 1X annexin-binding buffer (100uL) at a concentration of 1×10^6^ cells/mL. FITC Annexin V (5uL) and 100ug/mL PI working solution (1uL) were added to each 100 uL of cell suspension and incubated at room temperature for 15 minutes. After incubation 1X annexin-binding buffer (400uL) was added. For Caspase 3,7, cells were resuspended in RPMI media at a concentration of 1×10^6^ cells/mL. Cells were prepared for flow cytometry using the CellEvent™ Caspase-3/7 Green Flow Cytometry Assay Kit. Cell suspension (1mL) along with CellEvent™ Caspase-3/7 Green Detection reagent (1uL) were added to each flow cytometry tube and incubated for 30 minutes at 37C protected from light. In the final 5 minutes SYTOX™ ADDvanced™ Dead Cell Stain containing DMSO (1uL) was added.

For Violet Stain, cells were centrifuged for 5 minutes at 500g, supernatant was removed, and fixed by resuspension in 70% ethanol on ice for 1 hour. Cells were then centrifuged, supernatant removed and resuspended in 0.1% Triton X-100 + 1% BSA to PBS buffer with a concentration of 1×10^6^ cells/mL. Cells were prepared for flow cytometry using the FxCycle™ Violet Stain kit. Cell suspension (1mL) and FxCycle™ Violet stain (1uL) were added to each flow cytometry tube and incubated for 30 minutes at room temperature protected from light.

Flow analysis: After 48 hours of drug treatment cells were harvested and washed once in PBS. Samples were put on ice and brought to flow cytometer to be analyzed. After 2 additional days, cells were evaluated on a BD LSRFortessa cytometer. For all stains, the gating strategy was set to visualize side scatter area (SSC-A) vs forward scatter area (FSC-A) to distinguish melanoma cells from debris. To separate single cells from doublets, forward scatter area (FSC-A) vs forward scatter width (FSC-W) was plotted, except for the Violet stain, where BV421-A vs BV421-W was plotted.

### Dataset

The processed dataset comprised 1,874 cell-centered QPI sequences. For each annotated cell, its center pixel coordinate was selected at reference timepoint T0. A single fixed 100 × 100-pixel crop centered on this T0-defined coordinate was then extracted from the five frames preceding T0, the T0 frame itself, and the five subsequent frames T0-5 through T0+5. Each sequence was therefore stored as an 11-frame, 100 × 100-pixel, 16-bit TIFF stack. Each raw stack was loaded as a floating-point array with shape (T,H,W), where T=11 and H=W=100. Training experiments used global min–max intensity scaling by dividing each pixel by 65525. Thus, input values represented the original 16-bit QPI intensity scale mapped approximately to [0,1], while retaining the relative intensity relationships among timepoints within a sequence. For bootstrapped analysis of performance, the dataset was split randomly into train (1,311), validation (281), and test (282) samples, with evaluation reported on the held-out test set across 10 independent random seeds per configuration. All post hoc model evaluation and quantitative saliency analyses reconstructed the exact test partition from the saved configuration file associated with the analyzed run.

### Data augmentation

Training sequences received stochastic augmentation consisting of: random horizontal flip with probability 0.5, random vertical flip with probability 0.5, random 90°, 180°, or 270° rotation with probability 0.5, additive Gaussian noise with standard deviation 0.02 and probability 0.3, and cutout with probability 0.3, masking a square region with side length equal to 10% of the image dimension. Geometric transformations and cutout were applied to the full stack to preserve spatial alignment among frames. No augmentations were applied to validation or test data.

### 2D CNN models

2D models (EfficientNet-B0, EfficientNet-V2-S, ResNet-18, ResNet-34, ResNet-50, ResNet-101, MobileNet-V3-Small) used torchvision (github.com/pytorch/vision) pretrained CNN backbones adapted for the selected number of grayscale QPI frames as input channels. The first convolution was adapted to the input-channel count. For a single QPI frame, pretrained RGB kernels were averaged over the original color-channel dimension. For three-frame inputs, pretrained RGB kernels were retained. For inputs with more than three channels, pretrained kernels were repeated across channels and scaled by the number of repeats. The final classifier was replaced by a six-class linear layer.

### 3D CNN model

The 3D (temporal) model was based on the torchvision R(2+1)D-18 architecture (29). The first convolutional layer was modified to accept a single grayscale channel rather than three RGB channels. Its weights were initialized by averaging the pretrained RGB kernels across the input-channel dimension. The original classifier was replaced with a dropout layer (probability 0.3) followed by a six-output linear layer.

### Model training

Each model was trained for 100 epochs with batch size 32. To mitigate class imbalance, training data were sampled with replacement using a weighted random sampler. Models were optimized with AdamW using a learning rate of 10^-3^ and weight decay of 10^-3^. A cosine-annealing learning-rate scheduler was applied over 100 epochs. Training used focal loss,

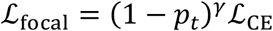

with focusing parameter γ = 2, where *p_t_* is the probability assigned to the true class and *ℒ*_CE_ is cross-entropy loss (30). At the end of every epoch, the model was evaluated on the validation partition. The checkpoint with the highest validation macro-F1 score was retained. The best performing model checkpoint was then evaluated on its held-out test split.

### Occlusion sensitivity analysis

We performed quantitative adaptive occlusion sensitivity analysis (AOSA) (31) on the test split using simple fixed-cuboid masks (20×20 pixel spatial patch, temporal crop size = 1, stride = 10) and the predicted class as the target. The raw effect at every spatiotemporal voxel is the drop in the target logit when that voxel is occluded, so positive values indicate regions whose occlusion lowers the model’s confidence in the predicted class.

### Statistics

Accuracies are given as mean +/-SD across all 10 bootstrapped runs.

## Results

Quantitative phase imaging results in live quantitative images suitable for morphologic profiling. Building on our prior success using QPI for cell state classification (5,7,17,32), we reasoned that incorporating time-series data would improve our ability to resolve subtle morphological differences among cell states with QPI. To generate distinct cell states for testing our deep-learning classification framework, we cultured healthy cells alongside cells exposed to five well-characterized drug treatments that induce apoptosis, ferroptosis, necroptosis, and growth arrest (Figure 1a). Flow cytometry-based validation using established markers of cell cycle and cell death pathway activation confirmed that the drug treatments resulted in the expected cell state (Figure S1). To dynamically capture the events associated with each cell state, we defined the event T0 critical frame between phenotypic changes associated with mitosis or cell death. For example, T0 for mitosis is defined as the last frame with a rounded cell just prior to splitting into two cells, while apoptosis T0 is defined by condensation, just prior to any blebbing or detachment. By selecting the centroid X,Y location for each cell at T0, we defined each event without requiring segmentation as the 100×100 pixel x N frame region around that centroid for 1874 total events across all classes.

**Figure 1.**
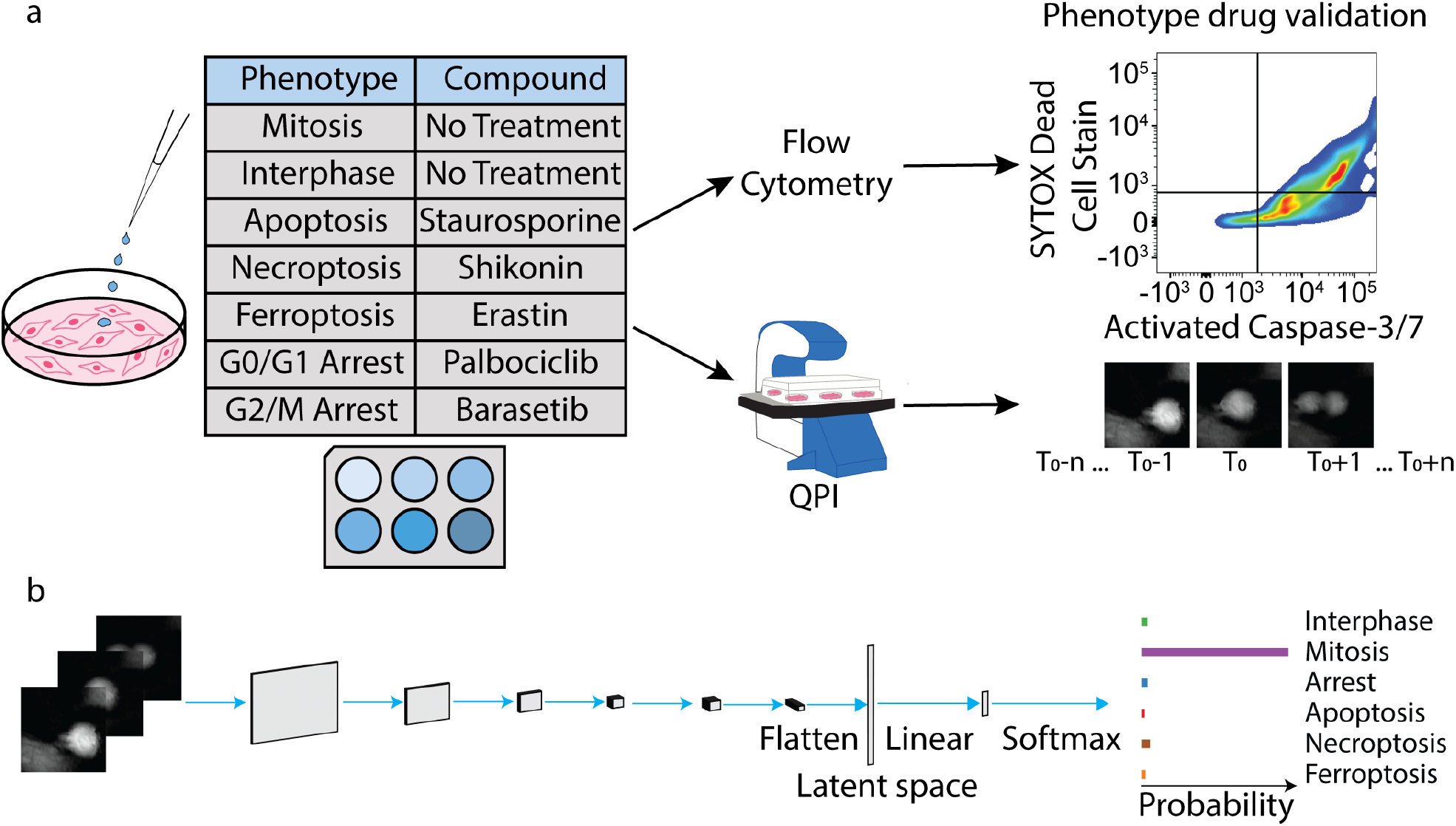
A deep learning pipeline classifies cell state based on dynamic imaging. (a) Imaging and annotation pipeline for phenotype classification. (b) Classifier training pipeline taking QPI data as input to predict cell phenotype.

Because cell imaging data contain spatial information across image slices, we considered both two-dimensional (2D) and three-dimensional (3D) convolutional neural network architectures (Figure 1b). A 2D CNN processes volumetric imaging data one slice at a time, extracting features from individual image planes without explicitly modeling relationships between neighboring slices. This approach is more computationally efficient and requires less memory, but it may lose important inter-slice spatial context. In contrast, a 3D CNN analyzes the full image volume simultaneously, allowing the network to learn volumetric features and spatial relationships across adjacent slices. As a result, 3D CNNs can provide richer feature representations and often achieve higher classification accuracy than 2D approaches by leveraging inter-slice information. For example, 3D CNNs have been shown to outperform 2D CNNs in ultrasound image classification by more effectively capturing spatial relationships across multiple slices (33). However, these benefits come with increased computational complexity, memory requirements, and training time.

Within the 3-frame time series signatures, dynamic differences distinguish different cell states (Figure 2a). Healthy cells are adherent with an intact cell membrane and nucleus. During mitosis, these cycling cells accumulate mass, round up, and divide into two round cells which regain the adherent morphology in 2-3 frames after division. In apoptosis, the most common form of programmed cell death, cells shrink, the cytoplasm and nucleus condense, and the cytoplasm fractures into distinctive apoptotic bodies (16,34). Necroptotic cells swell and the plasma membrane fractures, leading to translucent appearing cytoplasm and leakage of cell contents (35). Cells undergoing ferroptosis - cell death caused by iron-dependent production of lethal lipid peroxidation - conserve their structural integrity better than other dying cells, generally maintaining their adherent shape as the cytoplasm breaks down (36).

**Figure 2:**
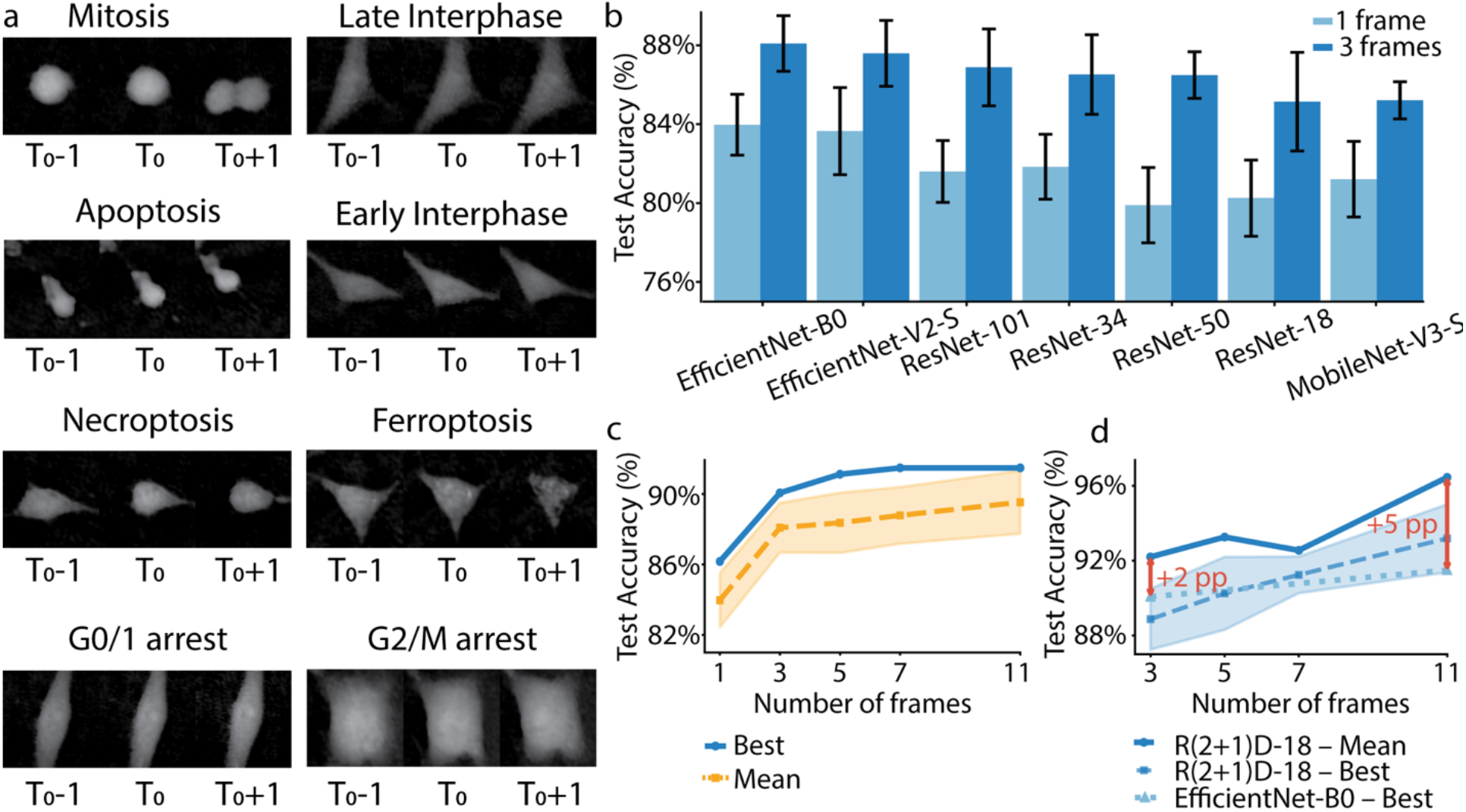
Incorporation of multiple frames improves overall accuracy of the deep learning classifier of cell state across multiple different model architectures. (a) 3 frame snapshots from different classes highlight differences in morphology characteristic of each cell state. (b) Mean test accuracy of seven 2D CNN architectures trained on single-frame and 3-frame inputs. Error bars show standard deviation of accuracies from 10 randomized training runs. (c) Mean and best test accuracies of EfficientNet-B0 across input window sizes of 1 to 11 frames. Yellow shaded region shows the standard deviation of accuracies from 10 randomized training runs. (d) Best-run test accuracy of EfficientNet-B0 (dotted line) and R(2+1)D-18 (solid line), and mean accuracy of R(2+1) D-18 (dashed line), across input window sizes of 3 to 11 frames. Blue shaded region shows standard deviation of accuracies from 10 randomized training runs.

To compare single frame evaluation to time series data, we considered both 2D and 3D architectures for our machine learning pipeline. To select a 2D CNN backbone, seven architectures were benchmarked at 1 frame (single middle frame, static morphology) and 3 frames (middle frame ± 1, minimal temporal context): EfficientNet-B0, EfficientNet-V2-S, ResNet-18, ResNet-34, ResNet-50, ResNet-101, and MobileNet-V3-Small (Figure 2b). Using only the middle frame, all architectures achieved moderate classification performance, reflecting the inherent difficulty of differentiating cell states from static morphology alone. EfficientNet-V2-S achieved the highest single-run accuracy at 87.23% (mean 83.7% ± 2.2%), followed by EfficientNet-B0 at 86.2% (mean 84.0% ± 1.5%), ResNet-50 at 84.8% (mean 81.8% ± 1.7%), ResNet-101 at 84.4% (mean 81.6% ± 1.6%), MobileNet-V3-Small at 83.7% (mean 81.2% ± 1.9%), and ResNet-18 and ResNet-34 both at 82.6% (means 80.2% ± 1.9% and 79.9% ± 1.9%, respectively). Expanding the inputs to 3 frames produced consistent improvements across all architectures, confirming the benefit of incorporating temporal dynamics into the classification pipeline. We further find improved overall classification with increased number of frames included in the data, up to incorporation of 11 frames (Figure 2c). The 1 to 3 frame transition yielded the largest single gain in accuracy, suggesting that even minimal temporal context dramatically improves class discriminability over static morphology (Figure 2c). Furthermore, when considering data that contains up to 11 frames, we find that using a 3D architecture shows improvements over 2D architectures (Figure 2d). From these data, EfficientNet-B0 was selected as the best-performing 2D architecture based on its superior accuracy and lower variance, indicating more reliable and reproducible training behavior. This consistent performance is critical for practical deployment and further temporal scaling experiments.

To assess whether a dedicated spatiotemporal architecture could surpass the best 2D model, EfficientNet-B0 was compared against R(2+1)D-18, which is a 3D CNN that applies spatiotemporal convolutions jointly across the frame and spatial dimensions at equivalent frame counts (29)(Figure 2d). Unlike EfficientNet-B0, R(2+1)D-18 continued to benefit substantially from additional frames beyond 3, with an 11-frame best accuracy of 96.5% representing a 4.2% improvement over its 3-frame performance. At 3 frames, R(2+1)D-18 (92.2%) outperformed EfficientNet-B0 (90.1%). The performance advantage of R(2+1)D-18 widened considerably at 11 frames: 96.5% versus 91.5%, with 10 misclassifications (3.5% error rate) compared to 24 (8.5%) for EfficientNet-B0. This growing gap with more frames indicates that 3D spatiotemporal convolutions scale far more effectively with increasing temporal context than 2D channel concatenation, and that the marginal saturation observed in EfficientNet-B0 reflects an architectural limitation in classification of dynamic events rather than an absence of useful temporal information in the QPI sequences.

Per-class test accuracy reveals that improvements in overall accuracy are distributed unequally across the six classes, and that the degree to which each class benefits from additional frames is closely tied to the nature of its temporal signature in QPI (Figures 3, S2). Mitosis is the best-performing class across all frame counts and the only one to show no meaningful accuracy gain with more frames (EfficientNet-B0: 96.9% at 1 frame, 96.0% at 11 frames; R(2+1)D-18: 97.1% at 3 frames, 98.9% at 11 frames). Mitotic cells adopt a distinctive rounded morphology with loss of substrate adhesion that produces unambiguous QPI signatures even in a single frame, so temporal context provides negligible additional discriminative power. Ferroptosis, by contrast, records the largest temporal gain of any class for EfficientNet-B0 (76.2% at 1 frame to 92.7% at 11 frames), and begins from the lowest single-frame baseline among all six classes. Ferroptosis lacks the sharply defined static morphological hallmarks of apoptosis or necroptosis. In a single QPI frame, ferroptotic cells resemble stressed interphase cells. Temporal sequences, however, expose characteristic, progressive accumulation of membrane irregularities and gradual decrease in density characteristic of this pathway, enabling substantially improved discrimination as more frames are included.

**Figure 3.**
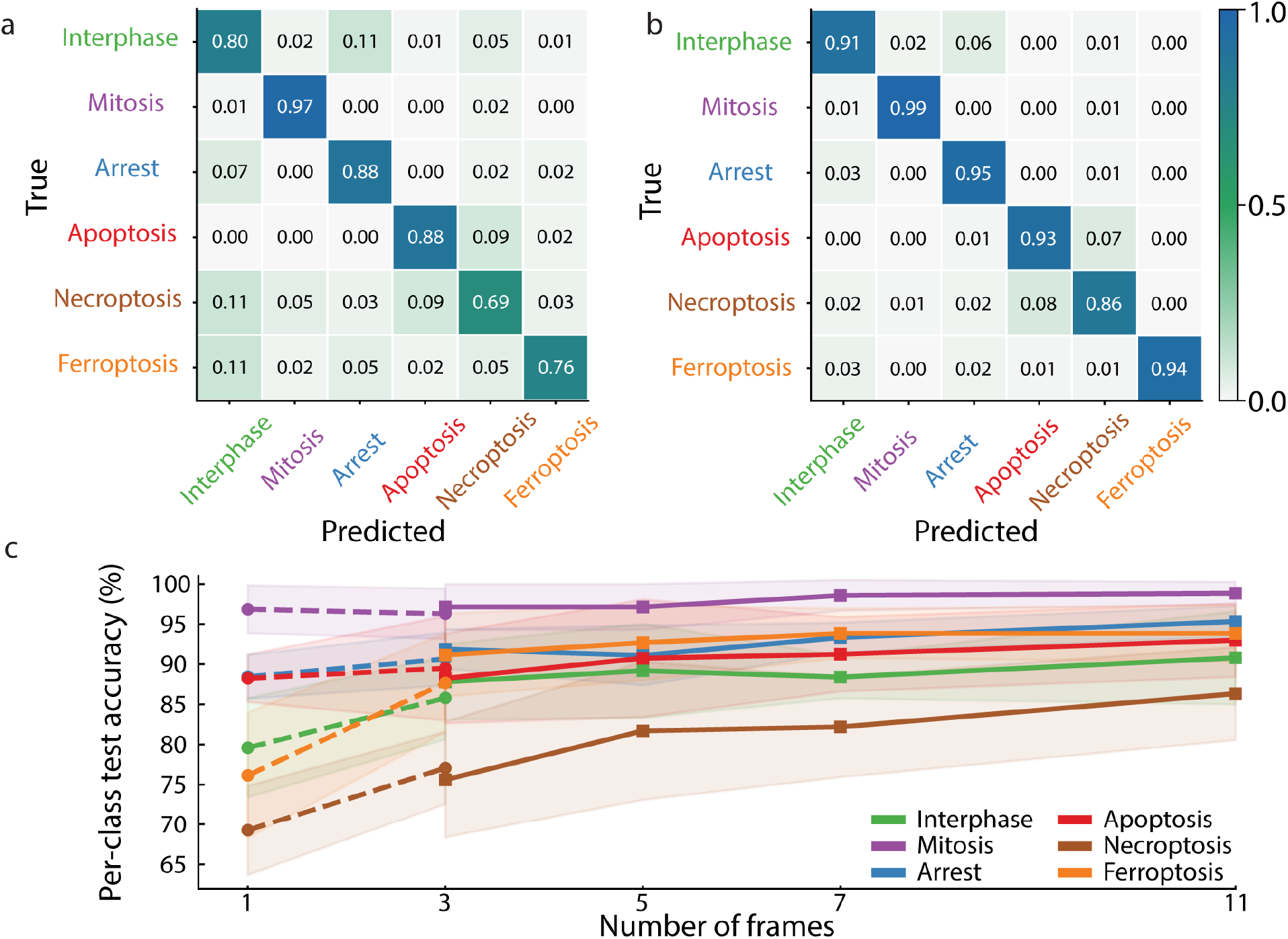
Incorporating multi-frame data into CNN models reduces misclassification errors of dynamic phenotypes. (a) Test dataset confusion matrix of EfficientNet-B0 trained with single frames. (b) Test dataset confusion matrix of R(2+1)D trained with 11 frames. Values in both confusion matrices are averages from 10 randomized training runs. (c) Per-class test accuracy for EfficientNet-B0 (dashed lines) and R(2+1)D-18 (solid lines) across all evaluated frame counts. Shaded regions show standard deviation of accuracies from 10 randomized training runs.

Necroptosis produces transient QPI signatures of a rapid phase density spike followed by abrupt collapse over the time window and requires multiple sequential frames to capture reliably, reflected by large gains with increasing temporal context (EfficientNet-B0: +9.3 percentage points, pp; R(2+1)D-18: +10.7 pp from 3 to 11 frames). In contrast, apoptosis shows modest but consistent improvement with temporal context (EfficientNet-B0: +2.8 pp; R(2+1)D-18: +4.8 pp from 3 to 11 frames), possibly due to relatively distinctive static markers of increasing density.

Cell cycle arrest and interphase are a frequently misclassified pair (Figure 3a-b), as arrested cells often maintain normal interphase morphology. Their discrimination relies on kinetic differences: actively cycling interphase cells exhibit slow phase density fluctuations reflecting ongoing biosynthesis and cytoskeletal dynamics, whereas arrested cells are comparatively static over the same temporal window. Consistent with this, both classes improve progressively with additional frames, with interphase showing the larger gain under EfficientNet-B0 (+7.6 pp vs. +3.3 pp for arrest). The residual class-level gap between arrest and interphase observed likely reflects the inherent difficulty of separating slow-cycling interphase cells from growth-arrested cells without molecular proliferation markers.

Improvements in classifier performance reflect latent space encodings in the classifier itself. We viewed latent space encodings using a UMAP, projected onto a consistent latent space using AlignedUMAP (37)(Figure 4a). Trajectories from 1 to 11-frame EfficientnetB0 models show a distinct improvement in clustering, with longer time intervals associated with tighter clustering (Figure 4b), a trend also captured in the 3D, R(2+1)D model (Figure 4c). These data indicate that unique temporal features in the QPI dataset are being captured by the model, which drives the observed improvement in performance (Figure 3).

**Figure 4:**
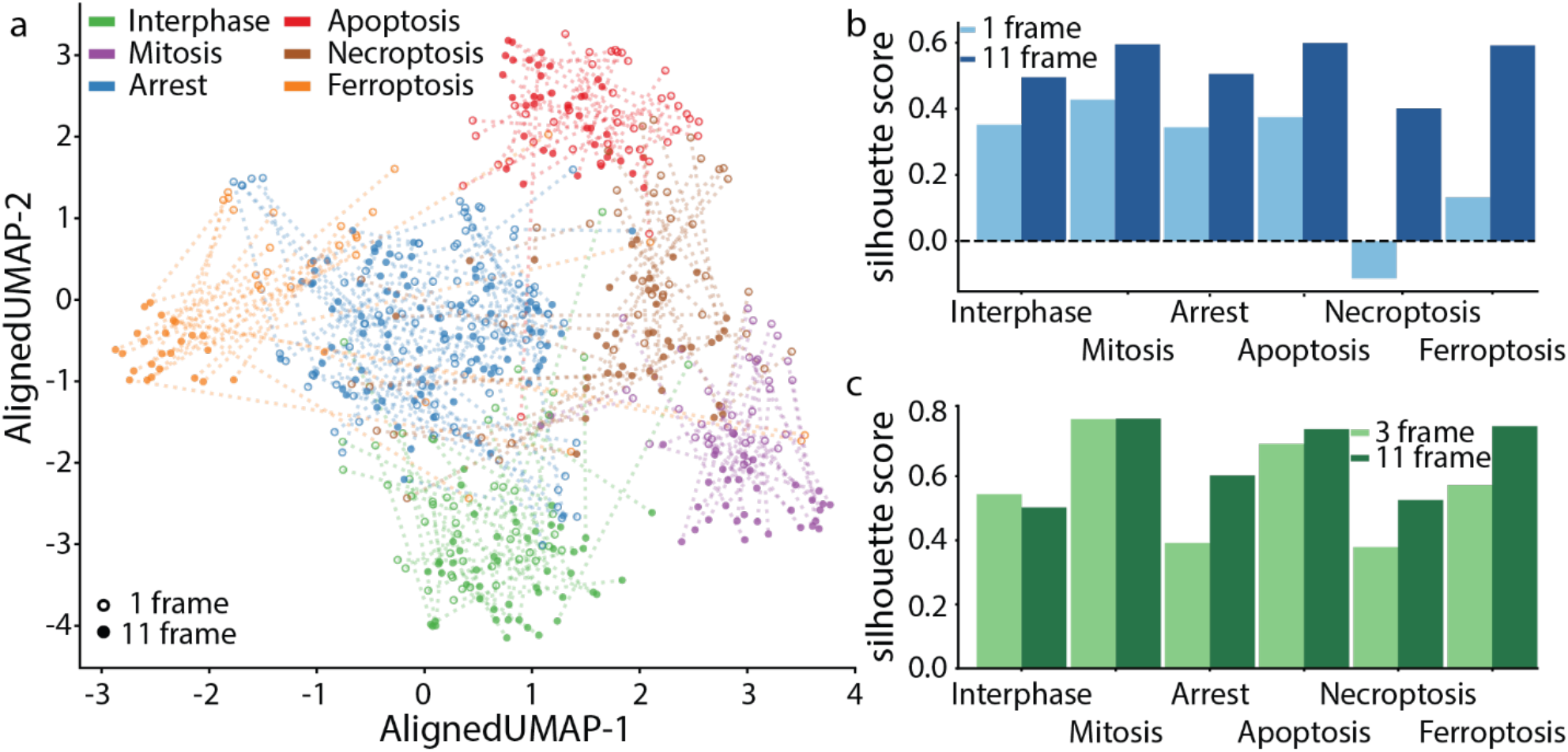
Features encoded in latent space show greater separation with more frames used in the classifier. (a) UMAP projections of latent features for 1 (open points) and 11 (closed points) frame EfficientnetB0. Color indicates cell state and matched open and closed points are connected (dotted line). (b) Silhouette scores of UMAP projections of latent features for 1 and 11 frame EfficientnetB0. (c) Silhouette scores of UMAP projections of latent features for 3 and 11 frame R(2+1)D.

To determine which portions of the temporal input contributed most strongly to R(2+1)D-18 classification, we applied adaptive occlusion sensitivity analysis (AOSA) to the held-out test split. For each class, AOSA scores were summarized across the 11-frame event-centered sequence by calculating the mean score of the top 15% highest-scoring pixels in each frame. The resulting frame-wise occlusion profiles showed that informative temporal regions were not uniformly distributed across the sequence but instead differed by cell state (Figure 5a).

**Figure 5.**
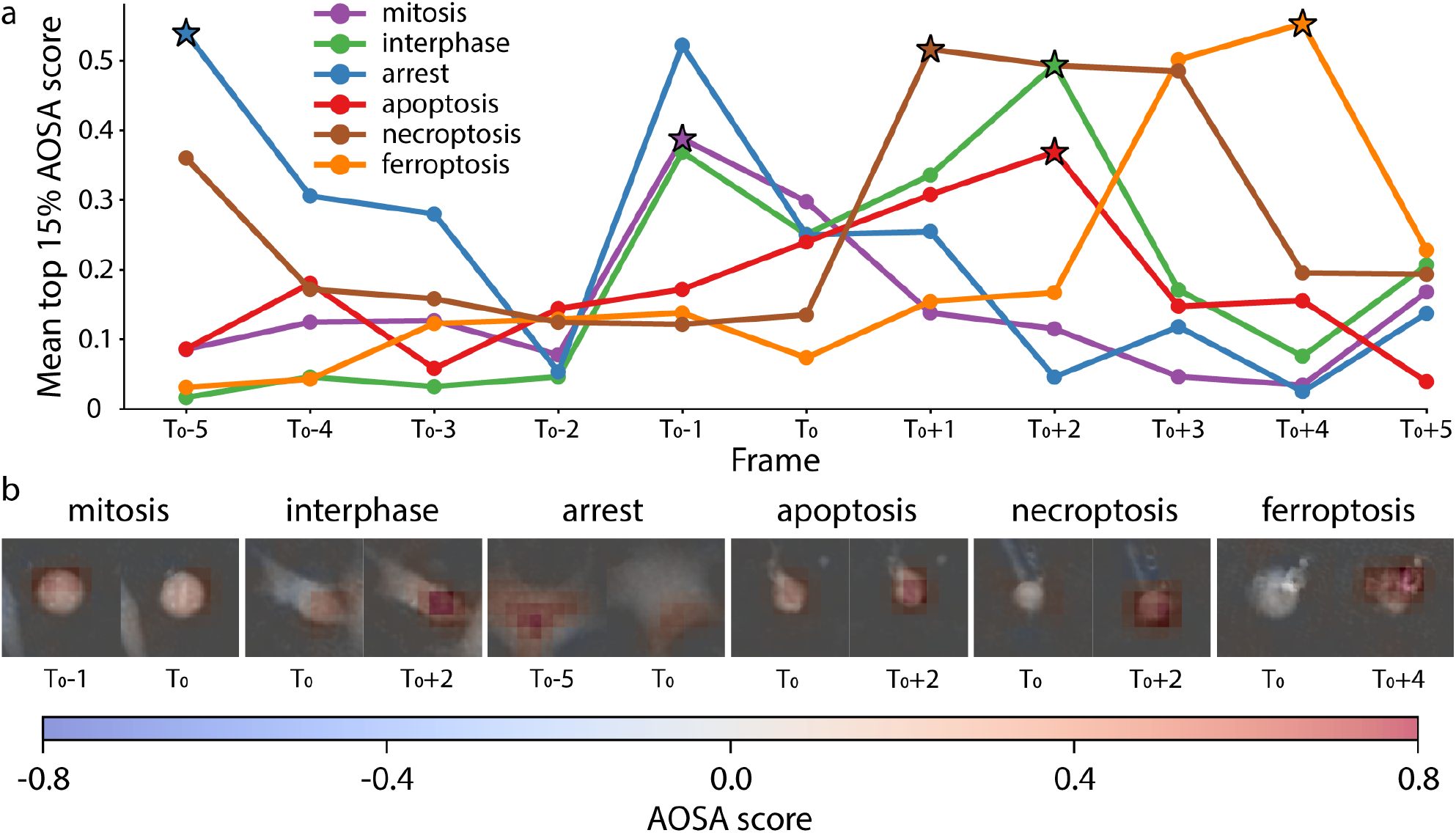
Adaptive occlusion sensitivity analysis identifies dynamic QPI features that contribute to cell-state classification. (a) Mean AOSA score for the top 15% highest-scoring pixels in each frame of representative 11-frame QPI sequences selected from each cell-state class. Scores are plotted across the event-centered temporal window from T–5 to T+5. Star markers indicate the frame with the highest mean top-15% AOSA score for each class, identifying the time point at which occlusion had the greatest effect on model confidence. (b) AOSA maps overlaid on QPI images at T0 and at the highest-scoring frame identified in panel A for each class. Red regions indicate image areas where occlusion decreased confidence in the predicted class, suggesting positive evidence used by the model, whereas blue regions indicate areas where occlusion increased confidence or reduced evidence for the predicted class. These results show that the model uses class-specific spatial features and temporally localized morphological changes to distinguish cell states.

Apoptosis showed elevated occlusion effects across the mid-to-late portion of the sequence, with the strongest response at frame 8, corresponding to T+2, after increased effects from frames 4–7. Growth arrest exhibited multiple informative time points, including a strong early effect at frame 1 and a second peak near frame 5. Ferroptosis was dominated by late-frame contributions, reaching its maximum at frame 9, with a secondary peak at frame 8. Interphase showed the highest occlusion effect near frame 6, corresponding to T0, whereas frames 1–3 produced near-zero effects, suggesting that the earliest frames provided limited supportive evidence for this class. Mitosis showed a sharper temporal profile, with the strongest occlusion effect at frame 5. In contrast, necroptosis displayed a more distributed pattern, with modest maxima around frames 4–5 and relatively uniform effects across the remainder of the sequence.

Together, these class-specific AOSA profiles indicate that the R(2+1)D-18 model did not rely equally on all input frames. Instead, different cell states were associated with distinct temporal windows in which occlusion of localized QPI features most strongly altered model confidence. These results support the interpretation that dynamic morphological information, rather than static morphology alone, contributes to the improved performance of the temporal CNN classifier.

## Discussion

Encoding live QPI as a multi-frame input allowed our classifier to resolve cell states that single images could not reliably separate. The amount of temporal context required for accurate classification varied between cell states and reflected the timescale over which their cell morphology changed. Classification accuracy rose with added frames, with the largest single gain occurring between one and three frames and further improvement continuing out to eleven frames (Figure 2). A 3D spatiotemporal network, R(2+1)D-18, converted this added temporal context into accuracy more effectively than 2D channel concatenation, reducing the test error rate to 3.5% at eleven frames (Figures 2, 3). Overall, these results indicate that dynamic information, rather than static morphology alone, drives accurate discrimination of cell cycle arrest and the major regulated cell death pathways in QPI.

The per-class pattern of improvement indicates that each state needs temporal context that is determined by the timescale of its underlying biological process. Mitosis produces an unambiguous sequence of rounding and division that completes within tens of minutes (38), so a single frame already captures its distinctive morphology. Accordingly, mitosis reached the highest accuracy at every frame count and gained essentially nothing from additional frames (Figure 3). Ferroptosis started from the lowest single-frame accuracy and showed the largest temporal gain of any class, consistent with the gradual accumulation of membrane irregularity and QPI derived density changes during ferroptosis (39), which became distinctive only when viewed across many frames. Necroptosis and apoptosis fell between these limits and improved steadily as added frames captured the swelling and membrane rupture of necroptosis and the condensation and fragmentation of apoptosis (22,34)(Figure 3). Our results suggest that the amount of temporal context provided to the models should be matched to the dynamics of the cell state being classified. States that undergo gradual morphological changes, such as ferroptosis, benefit from integrating information across many frames, whereas transient and morphologically distinctive events, such as mitosis, can be accurately classified from a single frame. The learned feature space reinforced this interpretation. Low-dimensional projections of the latent features showed increasingly distinct separation between cells states as frame count increased, and silhouette scores quantifying cluster quality rose accordingly for both architectures (Figure 4). The classes that overlapped most at low frame counts were the same ones that gained the most accuracy with additional frames, indicating that temporal context does not simply sharpen decision boundaries but reorganizes how the network represents each state.

Intrinsically 2D and 3D architectures responded very differently to this added temporal information. EfficientNet-B0, which treats successive frames as stacked image channels, captured the early gain from the first few frames but saturated thereafter, whereas R(2+1)D-18, which applies convolutions jointly across time and space, kept improving well beyond three frames (Figure 2). This may reflect a limit on how channel concatenation represents motion rather than an absence of useful information in the longer sequences.

Several misclassifications persisted even with the full 11-frame temporal context given here, marking the limits of the current approach. The largest errors were in classifying growth-arrested cells from cycling interphase cells, possibly because arrested and slow-cycling interphase cells share similar, static morphology and differ mainly in slow phase-density fluctuations indicating cell activity. A smaller but notable confusion arose between necroptosis and apoptosis. However, some necroptosis-inducing treatments can engage low-level caspase activity, a subset of cells may adopt intermediate phenotypes, and determining whether this reflects genuine pathway crosstalk or partial apoptosis within the population will require orthogonal molecular validation. Finally, the present study examined a single melanoma line (624-Mel) and relied on manual selection of each cell centroid to define the analysis window, so generalization across cell types and QPI platforms remains to be established.

Extending this framework will focus on removing the manual steps that currently limit throughput. Automated single-cell segmentation and tracking would define event windows without user input and let the classifier operate continuously across an entire imaging field, turning the method into a scalable, label-free readout of cell fate. Because QPI is non-perturbing and well suited to long-duration live imaging (24,25), such a pipeline could follow how heterogeneous populations respond to treatment in real time, flag rare or emergent phenotypes such as drug-resistant subpopulations (40), and support functional precision oncology approaches that test patient material directly (9). By showing that even modest amounts of temporal information sharply improve label-free discrimination of cell cycle arrest and distinct death pathways, this work provides a practical route to richer dynamic phenotypic profiling with QPI.

## Acknowledgements

We thank the HSC Flow Cytometry Core Facility at the University of Utah, Salt Lake City, UT, for providing access to flow cytometry instrumentation. We additionally thank Peter Egelberg for providing a Holomonitor M4 (Phase Holographic Imaging, PHI) used for the study and technical support.

## Author Contributions

Conceptualization: R.L.J.-T., T.A.Z.; Data Curation: K.M., S.A.; Formal Analysis: R.L.J.-T, S.A.; Funding acquisition: R.L.J.-T., T.A.Z.; Investigation: K.M, A.D.; Methodology: S.A., R.G.Z, R.L.J.-T, T.A.Z.; Project Administration: R.G.Z, R.L.J.-T, T.A.Z.; Resources: R.L.J.-T., T.A.Z.; Validation: S.A., R.L.J-T.; Visualization: S.A., K.M., R.L.J-T.; Software: S.A.; Writing – original draft: S.A., K.M., R.G.Z.; Writing – review and editing: S.A., K.M., R.G.Z, R.L.J.-T., T.A.Z.; Supervision: R.L.J.-T., T.A.Z.

