## Supplementary material for "Temporal dynamics improves machine learning-based prediction of cell state from quantitative phase imaging": Supplemtal Material

**Contents:**

**Supplementary Figures 1-3**

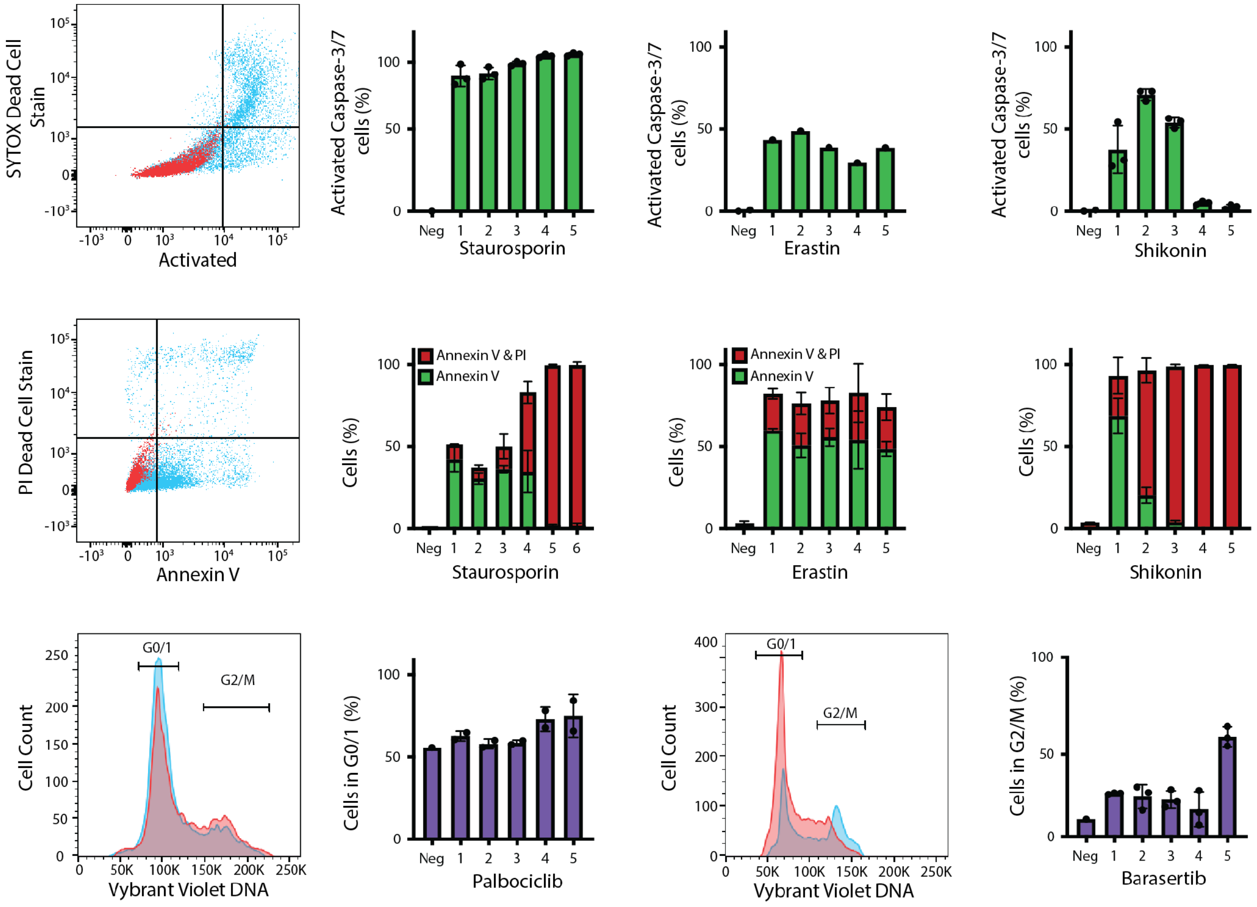


**Supplementary Figures S1. Flow cytometry validation of drug-induced cell states used for DHM classification.**

Flow cytometry was performed after 48 h of drug treatment to confirm that the small-molecule perturbations used for M4 HoloMonitor imaging induced the expected cell-death and cell-cycle phenotypes. Cells were treated with DMSO control or one of five drug conditions using the same treatment scheme used for dynamic QPI acquisition. Representative gating plots and summary quantification are shown for apoptosis/death-marker assays and cell-cycle analysis. Cells were initially gated by forward and side scatter to exclude debris, and single cells were identified using FSC-A/FSC-W gating or BV421-A/BV421-W gating for FxCycle Violet DNA-content analysis. Annexin V/propidium iodide and CellEvent caspase-3/7/SYTOX staining were used to quantify apoptotic and membrane-compromised cell populations across treatment conditions. FxCycle Violet staining was used to quantify DNA-content distributions, including G0/G1 and G2/M populations. Bar plots summarize marker-positive or gated populations across the negative control and drug-treated conditions, with replicate measurements shown. These flow cytometry results confirmed that the treatments produced distinct apoptotic, necroptotic, ferroptotic, and growth-arrest-associated phenotypes for downstream label-free DHM classification.

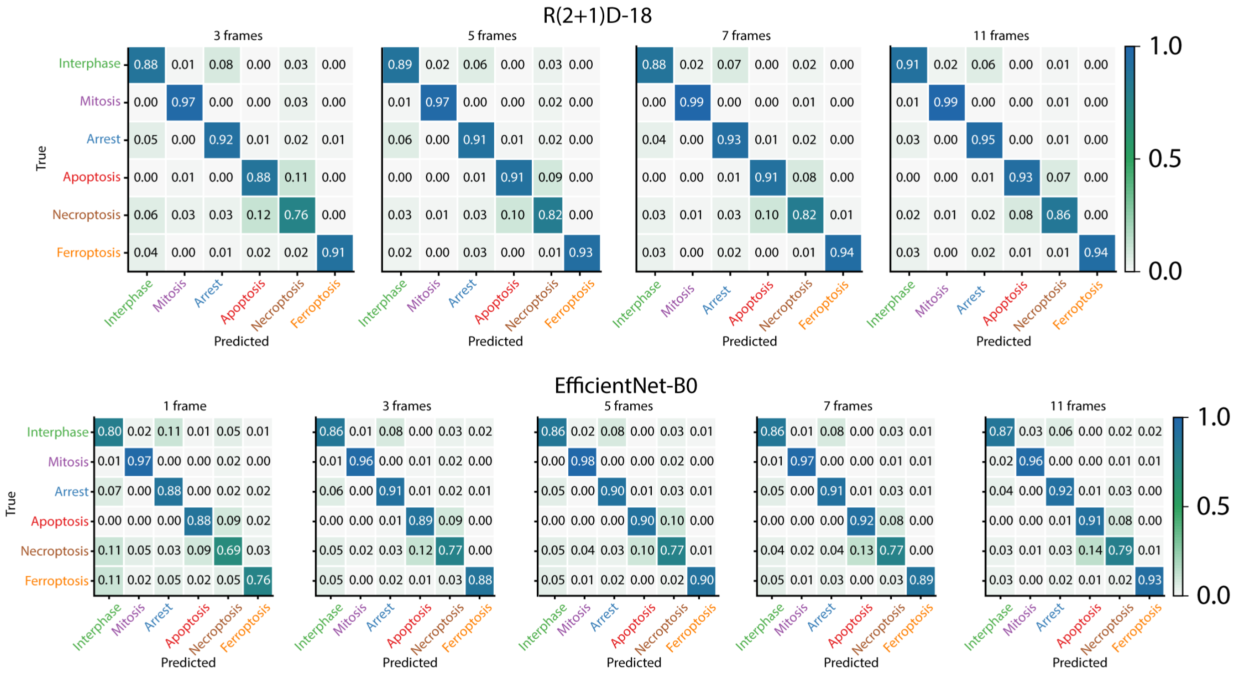


**Supplementary Figure S2. Confusion matrices for R(2+1)D-18 and EfficientNet-B0 across different temporal input lengths.**

Confusion matrices for R(2+1)D-18 (a) and EfficientNet-B0 (b) were generated from predictions on the held-out test set and averaged across 10 independently trained models for each input frame number. Rows indicate the true cell-state labels, and columns indicate the predicted labels. Matrix values represent the mean normalized prediction frequency across the 10 training runs. Stronger diagonal values indicate accurate classification, whereas off-diagonal values indicate recurring misclassification patterns between cell states. These matrices show how increasing the number of input frames affects class-level performance for the 3D temporal model, R(2+1)D-18, and the 2D CNN architecture, EfficientNet-B0.

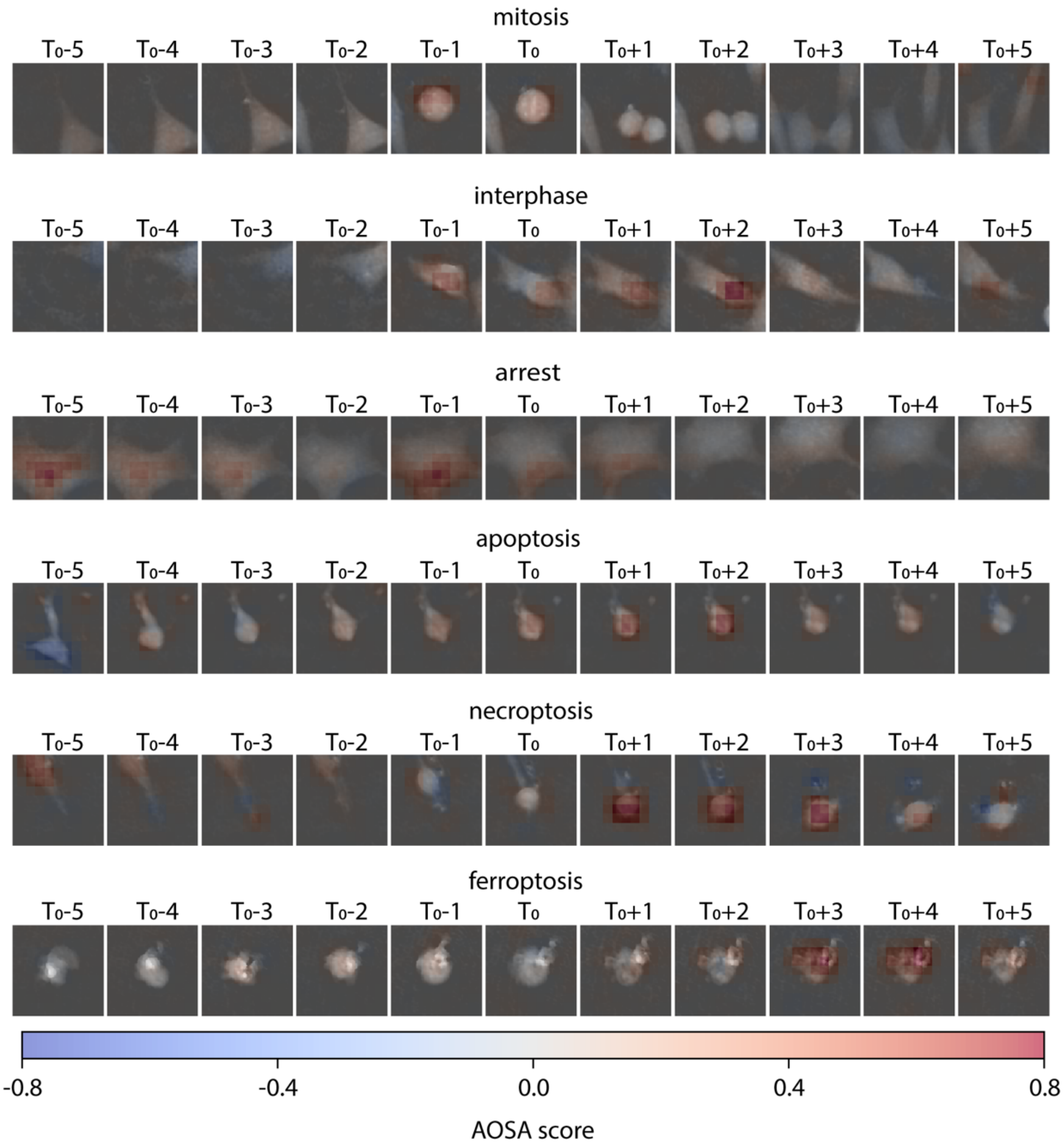


**Supplementary Figure S3. Adaptive occlusion sensitivity analysis maps overlaid on QPI image sequences for representative cell-state classes.**

Representative 11-frame quantitative phase imaging sequences are shown for mitosis, interphase, growth arrest, apoptosis, and necroptosis, spanning five frames before to five frames after the event-centered frame (T−5 to T+5). T0 denotes the frame used to temporally align each event sequence. Adaptive occlusion sensitivity analysis (AOSA) maps were generated using the best-performing R(2+1)D-18 model and overlaid on the corresponding QPI images to identify spatial and temporal regions that contributed most strongly to the model prediction. Positive AOSA scores, shown in red, indicate regions where occlusion reduced confidence in the predicted class, suggesting positive evidence for that class, whereas negative AOSA scores, shown in blue, indicate regions where occlusion increased confidence or reduced evidence for the predicted class. These overlays suggest that the model uses both localized morphology and dynamic changes across the event window to distinguish cell states.
